# A simple method to classify behaviours of aerial predators using tri-axial high-frequency accelerometers

**DOI:** 10.1101/2025.02.01.636019

**Authors:** D. García-Silveira, S. Benhamou, L. Lopez-Ricaurte, J. Hernández-Pliego, O. Duriez, J. Bustamante

**Author notes:** Currently as (D. García-Silveira).

## Abstract

**Background:** Remote monitoring of animals has opened new avenues in behavioural research. The advent of animal-borne accelerometers makes it possible to derive the body posture and activity level, and thereby to identify fine-scale behaviours with no need for direct observation. This is particularly useful for species that forage or hunt beyond human sight in sometimes hard-to-access or remote areas. However, the complexity of numerous methods developed for this purpose may discourage researchers from effectively using acceleration data.

**Results:** We present a simple and straightforward approach to identify and classify behaviours based on tri-axial high-frequency (25 Hz) acceleration data. This method relies on constructing a decision tree to assign predefined behaviours to time series of acceleration data using four accelerometer-based parameters that are biologically relevant and easy to interpret. Initially, we manually labelled sequences of acceleration data with the behaviours using paired high-frequency (i.e. 1 Hz) GPS tracking data as ground-truth. Then, threshold values of the acceleration-based parameters were objectively determined based on receiver operating characteristic (ROC) analysis. Finally, we built the decision tree to automatically classify the behaviours of the whole dataset and tested further improvements. As a case study, we used two falcon species (lesser kestrel *Falco naumanni* and common kestrel *F. tinnunculus*). Our simple method achieved an overall accuracy of 97.6% in classifying the following behaviours: perching (99.9%), soaring-gliding (99.3%), flapping (93.1%) and hovering (86.4%). We also showed that kestrels’ prey capture attempts can be identified from the tri-axial high-frequency accelerometers due to diagnostic signatures (the so-called ‘spikes’).

**Conclusions:** Our simple method has broad applicability to free-ranging aerial predators. We show that behavioural categories can be accurately classified from a relatively small subset of manually labelled acceleration data (43,075 seconds). Although quantifying hunting success needs further investigation, the detection of prey capture attempts opens new possibilities for studying the hunting behaviour of aerial predators.

## Introduction

Technological innovations in tracking systems have facilitated major advances in the study of animal movement by allowing researchers to overcome many issues associated with observational studies, such as observer effects, limited spatial scales or time-consuming fieldwork campaigns (Kays et al. 2015, López-López 2016). The ongoing miniaturisation and sophistication of tracking devices have broadened the range of species that can be studied with an unprecedented spatio-temporal resolution (Khan et al. 2016, Nathan et al. 2022). Simultaneously, the development of animal-borne sensors has provided remote access to behavioural, physiological and environmental variables (e.g. body acceleration, diving depth, heart rate, stomach temperature, wind speed, salinity), providing new insights into the ecological drivers that influence movement (Wilmers et al. 2015).

Tri-axial accelerometers are widely used sensors (Joo et al. 2022). By measuring acceleration in 3D at very high temporal resolutions (from 0.5 Hz to 10,000 Hz, Brown et al. 2013), these devices can determine the body posture as well as the activity level, which is a proxy of energy expenditure (Wilson et al. 2020). Three approaches can be used to translate the time series of raw acceleration data into specific behaviours (Shamoun-Baranes et al. 2012, Yu et al. 2021). (*i*) Direct classification based on expert opinion, with (Halsey et al. 2009) or without (Holland et al. 2009) ground-truth data from visual or video observations, requires predefined behaviours and is appropriate when acceleration signatures are easily identifiable (Yoda et al. 2004, Sur et al. 2023). (*ii*) Unsupervised machine learning or clustering identifies groups of common signatures in acceleration data and does not require any previous knowledge of behaviours or ground-truth data, but does need an interpretation of the identified clusters and may result in erroneous classifications when similar signals occur in different clusters (Watanabe et al. 2012, Chimienti et al. 2016). (*iii*) Supervised machine learning uses ground-truth data to train models for automatic classification and, while fast and reliable on predefined behaviours, it cannot detect unknown ones (Korpela et al. 2020).

Accelerometers have thus enabled the classification of behaviours from large datasets (e.g. ∼3.5 million daily values per axis at 40 Hz sampling, Fehlmann et al. 2017), becoming a well-established research tool to study time-activity budgets (Van Donk et al. 2020), foraging strategies (Williams et al. 2014) and biomechanics (Sato et al. 2009), but challenges remain in detecting and interpreting specific events. For instance, prey capture attempts by diving marine predators (e.g. Adélie penguins *Pygoscelis adeliae*, Watanabe and Takahashi 2013; Steller sea lions *Eumetopias jubatus*, Viviant et al. 2010) and by large terrestrial mammals (e.g. pumas *Puma concolor*, Wang et al. 2015; African leopards *Panthera pardus*, Wilmers et al. 2017; Canada lynxes *Lynx canadensis*, Studd et al. 2021) have been identified in combination with other sensors (e.g., time-depth recorders, acoustic recorders, GPS data). For other species, identifying diagnostic acceleration signatures offers new opportunities to detect such events, including prey caching in arctic foxes (*Vulpes lagopus*) or the distinct movement of Eurasian spoonbills (*Platalea leucorodia*) upon capturing a prey (Clermont et al. 2021, Lok et al. 2023).

In this study, we aimed at determining to which extent the behaviour and prey capture attempts by aerial predators such as lesser kestrels (*Falco naumanni*) and common kestrels (*F. tinnunculus*) could be identified using tri-axial high-frequency acceleration data alone. These two closely related falcons share flight modes (soaring-gliding and flapping flight) and hunting techniques (either by hovering flight or from a perch) (Village 1990, Fuchs et al. 2015, Hernández-Pliego et al. 2017b). We relied on paired acceleration data and high-frequency GPS tracking data to manually label time series of acceleration data with the predefined behaviours that we aimed at identifying (i.e. soaring-gliding, perching, flapping, hovering). Obviously, high-frequency GPS tracking can be performed only for short durations due to limited battery capacities. Furthermore, manual labelling is very time-consuming. We therefore investigated the extent to which behaviours identified using paired acceleration and GPS data could be accurately and reliably classified using acceleration data alone. For this purpose, we designed a decision tree involving four accelerometer-based parameters. We also attempted to identify the kestrels’ prey capture attempts by paying particular attention to specific acceleration signatures that were due to the abrupt movements involved (dives or strikes), also assessing the hunting method (perch-hunting or flight-hunting).

## Methods

### Study areas

Our study was conducted in two areas of south-western Spain where both species breed sympatrically (Martínez-Padilla 2016, Ortego 2016). In the first area, ‘La Palma’ (N 37.391145, W - 6.557886), the landscape is dominated by crops such as olives, wheat, sunflower and chickpeas, whereas the second area, ‘Doñana’ (N 37.070806, W -6.300587), is characterised by natural marshes and grasslands.

### Tracking

Kestrels were trapped using either bow-net traps or a remote-controlled sliding door at the nest box. We deployed bio-loggers combining high-frequency (1 Hz) GPS receptors and high-frequency (25 Hz) tri-axial accelerometers (Axy-Trek 3, TechnoSmArt Europe Srl., Rome, Italy) on 17 lesser kestrels (nine females and eight males) and four common kestrels (three females and one male). Individuals were tagged during the nestling periods of 2019 and 2020, from mid-May to early July, when both parents were provisioning their nestlings. The bio-loggers were protected with a heat-shrinking tube and attached to the kestrels using a backpack harness made of Teflon ribbon. The total equipment mass did not exceed 4% of lesser and common kestrels’ mean body mass (ca. 130 g and 200 g, respectively), and thus is within the accepted standards for animal welfare in research (Barron et al. 2010).

While accelerometers recorded data continuously, GPS receptors were programmed to acquire 3D location and instantaneous speed only during daylight hours (05:00-20:00 h) to save battery. The bio-loggers were programmed to start working 24 hours after their deployment to avoid monitoring potentially abnormal behaviour due to capture stress. As they could only store data on-board, we had to recapture the individuals to download the data. We collected data for 157 daylight hours (tracking period per individual: mean ± SD = 7.14 ± 2.87 hours). We removed the few GPS locations with low accuracy (i.e. computed from less than 4 satellites) and those that generated an inter-location ground speed higher than 50 km/h. Furthermore, tracked kestrels were monitored at nine nest boxes (seven occupied by lesser kestrels and two by common kestrels) using cameras (portable Ltl Acorn 5310 with an IR illuminator) that recorded videos of ten seconds when detecting movement. These video recordings were synchronised with the data collected by the bio-loggers.

### Acceleration data analyses

Call ***a*** *= (a*_*x*_, *a*_*y*_, *a*_*z*_) to the acceleration vector obtained at a given time, where *a*_*x*_, *a*_*y*_ and *a*_*z*_ correspond to the values obtained for the surge (tail-to-head), sway (right-to-left) and heave (belly-to-back) axes of the accelerometer, respectively (see Fig. 1 in Benhamou 2023). This vector combines the ‘static’ acceleration, which is due to gravity and reflects the body’s posture, and the ‘dynamic’ acceleration, which is due to changes of linear speed caused by the animal’s movement and reflects its activity (Shepard et al. 2008). The static component was estimated by the mean vector **ā** *= (ā* _*x*_, *ā* _*y*_, *ā* _*z*_) using a 1-s sliding window, and the dynamic body acceleration (hereafter DBA) was computed as the norm ||**d**|| of the vector difference **d** *= (d*_*x*_, *d*_*y*_, *d*_*z*_*) =* **a** *–* **ā**. We kept a single value for **ā** (through subsampling) and ||**d**|| (through averaging) per second, so the amount of data to process in subsequent analyses were reduced by 25-fold without a noticeable loss of information (Benhamou 2023). DBA was used as a proxy for energy expenditure (Qasem et al. 2012) and the static acceleration was used to compute the elevation angle (or longitudinal inclination) φ = sin^-1^(*–ā* _*x*_*/*||**ā** ||), which represents the vertical angular deviation of the surge axis with respect to the horizontal plane. An elevation angle close to zero means that the surge axis was horizontal and therefore the kestrel was laying down or flying, whereas, with the convention we used (a negative *ā* value obtained for a given axis means the axis was pointing upwards, i.e. in the opposite direction of gravity), a highly positive elevation angle means that the kestrel was tilted upwards and thus standing, and a negative elevation angle means that the kestrel pointed downward. We also computed, for every second, the number of times that *a*_*z*_ crossed the value –1*g* (i.e. –9.81 m/s^2^) and the coefficient of variation (hereafter CV) of the delay between two of such occurrences. A low CV indicates regular moves, which are due to wingbeat and whose frequency can be estimated as half the number of times that *a*_*z*_ crossed the value –1*g* (Fig. 1). Finally, we also designed a fifth acceleration parameter, which is a binary parameter that indicated the presence or absence of a distinctive high value of *a*_*z*_ within each 1-s interval (see below *Identification of prey capture attempts* for details).

**Figure 1.**
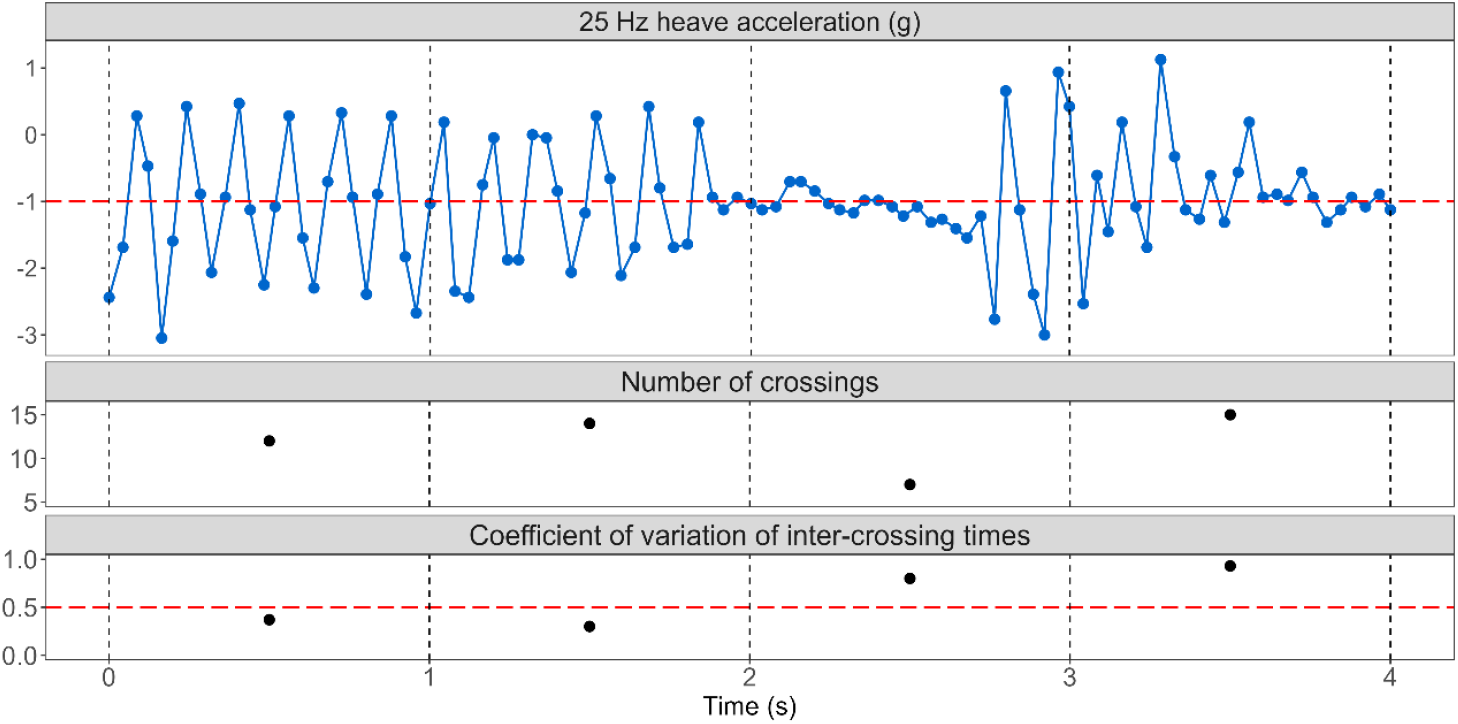
Example of a 4-s period that showed how regular moves in the heave axis (upper panel) resulted in high number of times that *a*_*z*_ crossed the value –1*g* and in low coefficients of variation (CV) of the delay between two of such occurrences (first two seconds). Irregular moves can be identified by high CV (last two seconds).

### Identification of behaviours based on paired acceleration and GPS data

We aimed at identifying four behaviours: perching, soaring-gliding, flapping and hovering (Fig. 2). Perching is characterised by low DBA, positive elevation angle and, when available, null GPS instantaneous speed (Fig. 2A). Soaring-gliding flight is characterised by low DBA, slightly negative elevation angle and, when available, GPS instantaneous speed > 1 km/h (Fig. 2B). Flapping and hovering flights differed from soaring-gliding in showing higher DBA and, as they involved regular oscillations in *a*_*z*_, lower CV (Fig. 2B-D). Flapping and hovering can be distinguished based on wingbeat frequency (usually higher in hovering) (Fig. 2C-D) and, when available, the GPS instantaneous speed (> 10 km/h vs. ca. 0 km/h, respectively). Sometimes, the distinction between the different types of flight was blurred for a few seconds (see Fig. 1, last two seconds). Those time sequences were characterised by high values in CV and DBA and were labelled as ‘irregular flight’. Although this is not a proper behaviour, and thus was not included when assessing the accuracy and precision of the classification model (Table 1), we used these irregular flight intervals to select the threshold in CV to identify flapping and hovering (Fig. 4C). As all the predefined behaviours could be easily characterised based on paired acceleration and GPS data, we manually labelled time sequences of each behaviour following the approach of direct classification (see Fig. S1 to explore the interactive web app created for that purpose). The minimum sample unit to label behaviours was 1-s interval. We then tested to which extent the behaviours could be accurately identified using acceleration data alone.

**Table 1.**
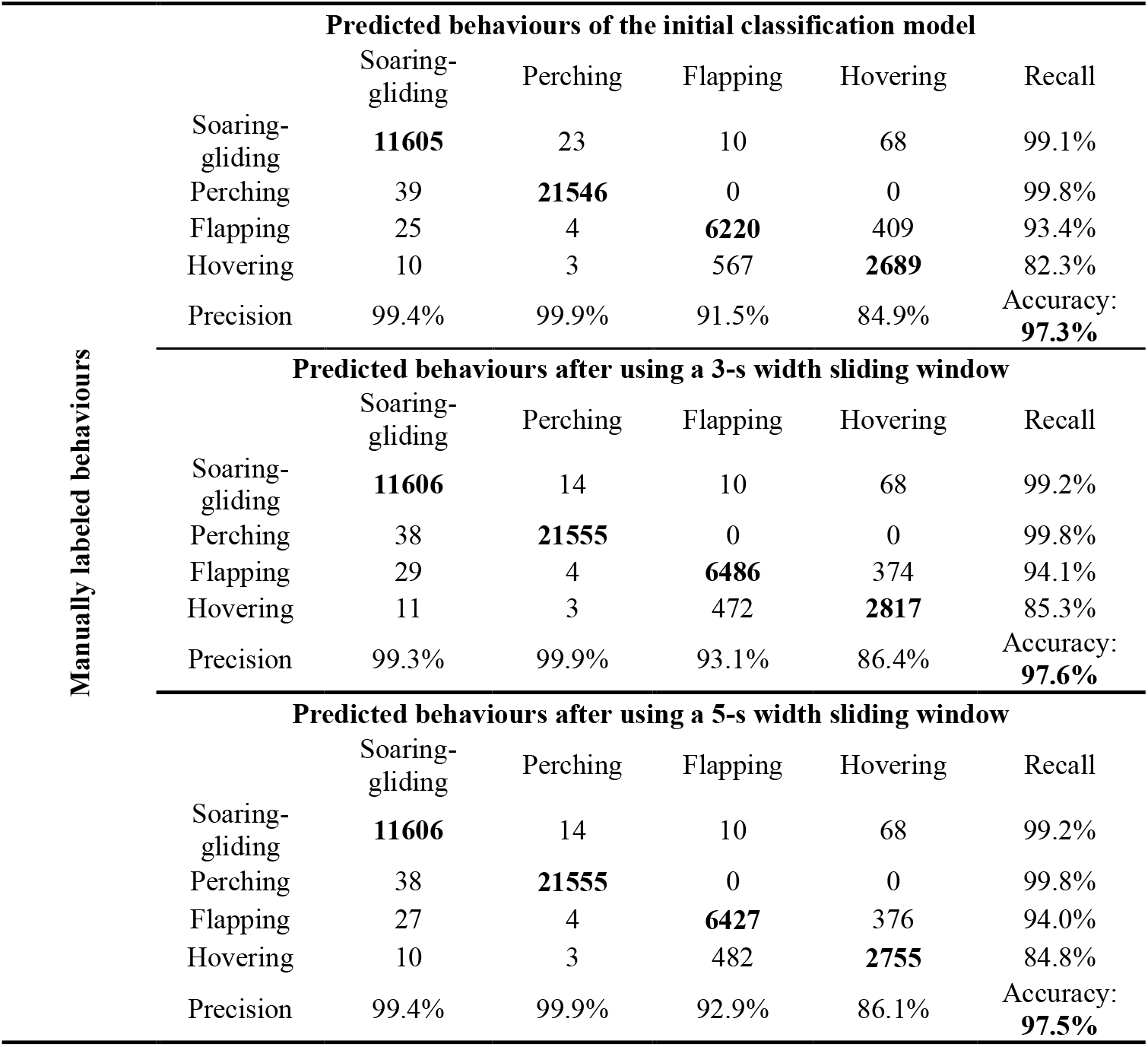
Confusion matrices for the initial classification model and the model with the 3-s width sliding window. Observations correctly classified (true positives) are shown in bold. The precision = true positives/(true positives + false positives) and the recall (or sensitivity) = true positives/(true positives + false negatives) are indicated for each behaviour. The accuracy = (true positives + true negatives)/(true positives + true negatives + false positives + false negatives) makes it possible to evaluate the classification globally.

**Figure 2.**
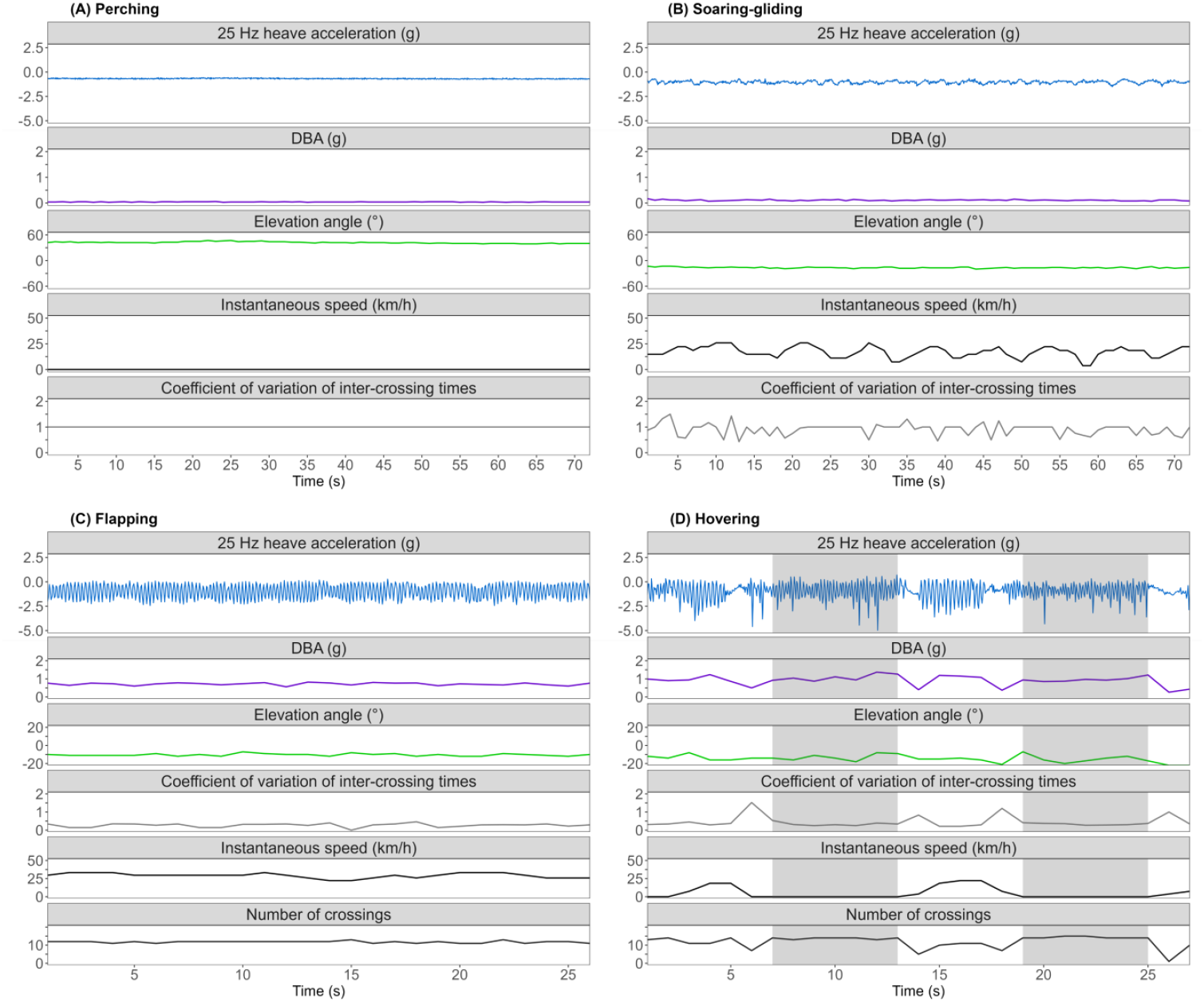
Parametrisation of kestrels’ behaviours. Example of time sequences that were manually labelled as soaring-gliding, perching, flapping and hovering. Grey areas in panel D indicated the hovering events, distinguished from flapping by zero GPS instantaneous speed and higher values in the number of times that heave acceleration crosses –1*g*.

### Classification, model fitting and validation

The decision tree was based on four acceleration parameters (DBA, elevation angle, number of crossings and CV of inter-crossing times) (Fig. 3). The threshold values were derived from a receiver operating characteristic (ROC) analysis (Fig. 4) based on 43,075 1-s intervals of manually labelled data: 21,601 s of perching from 20 individuals, 11,751 s of soaring-gliding from 21 individuals, 7,027 s of flapping from 21 individuals, 2,428 s of hovering from 19 individuals and 268 s of irregular flight from 20 individuals. As the mean value per individual for every parameter involved in the decision tree did not differ significantly between the two species (Kruskal-Wallis test: *P* >0.05 in all cases), we designed a single decision tree that could be applied to both species.

**Figure 3.**
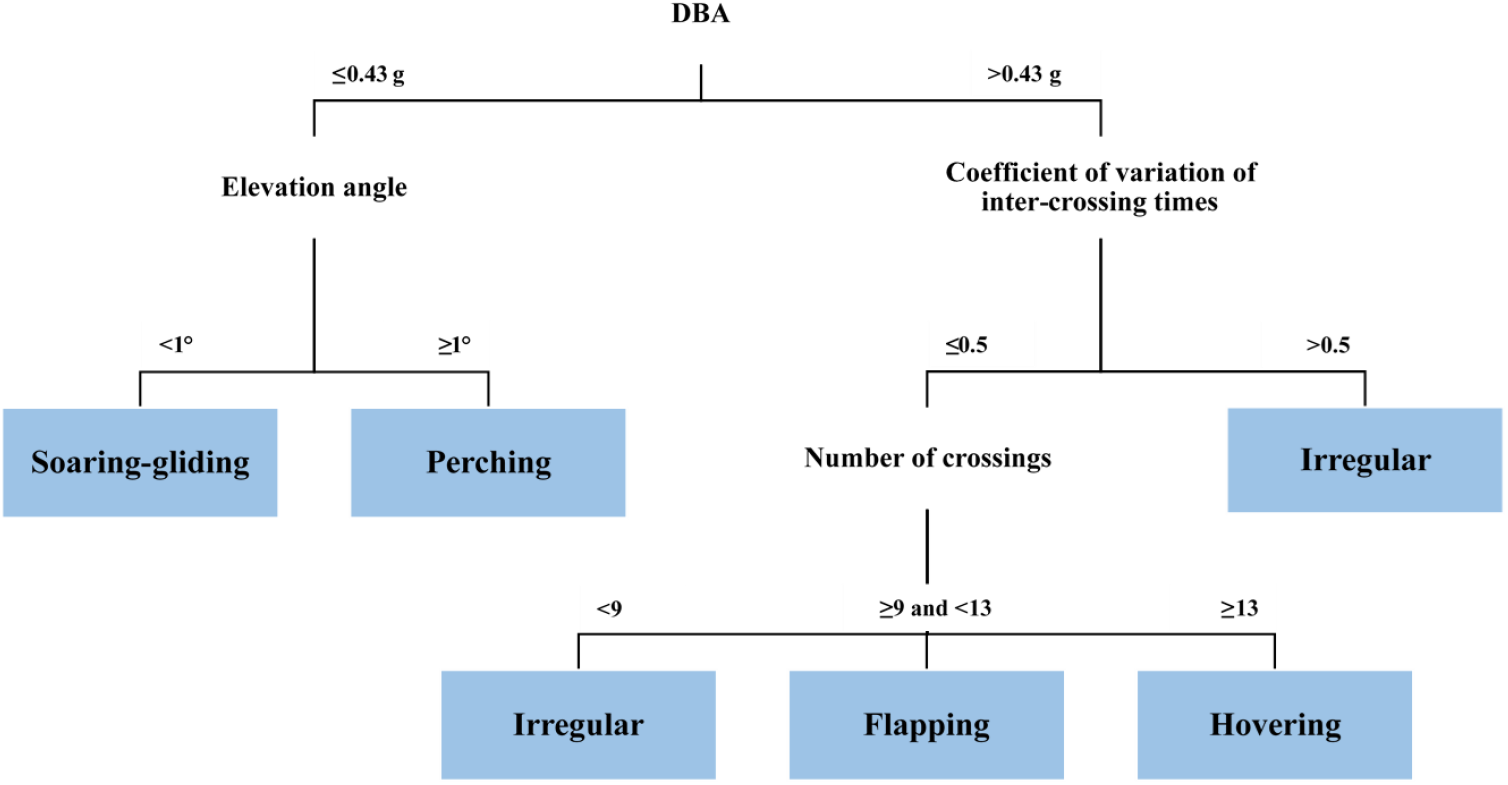
Decision tree and threshold values used to automatically classify the behaviours of kestrels based on four accelerometer-based parameters.

**Figure 4.**
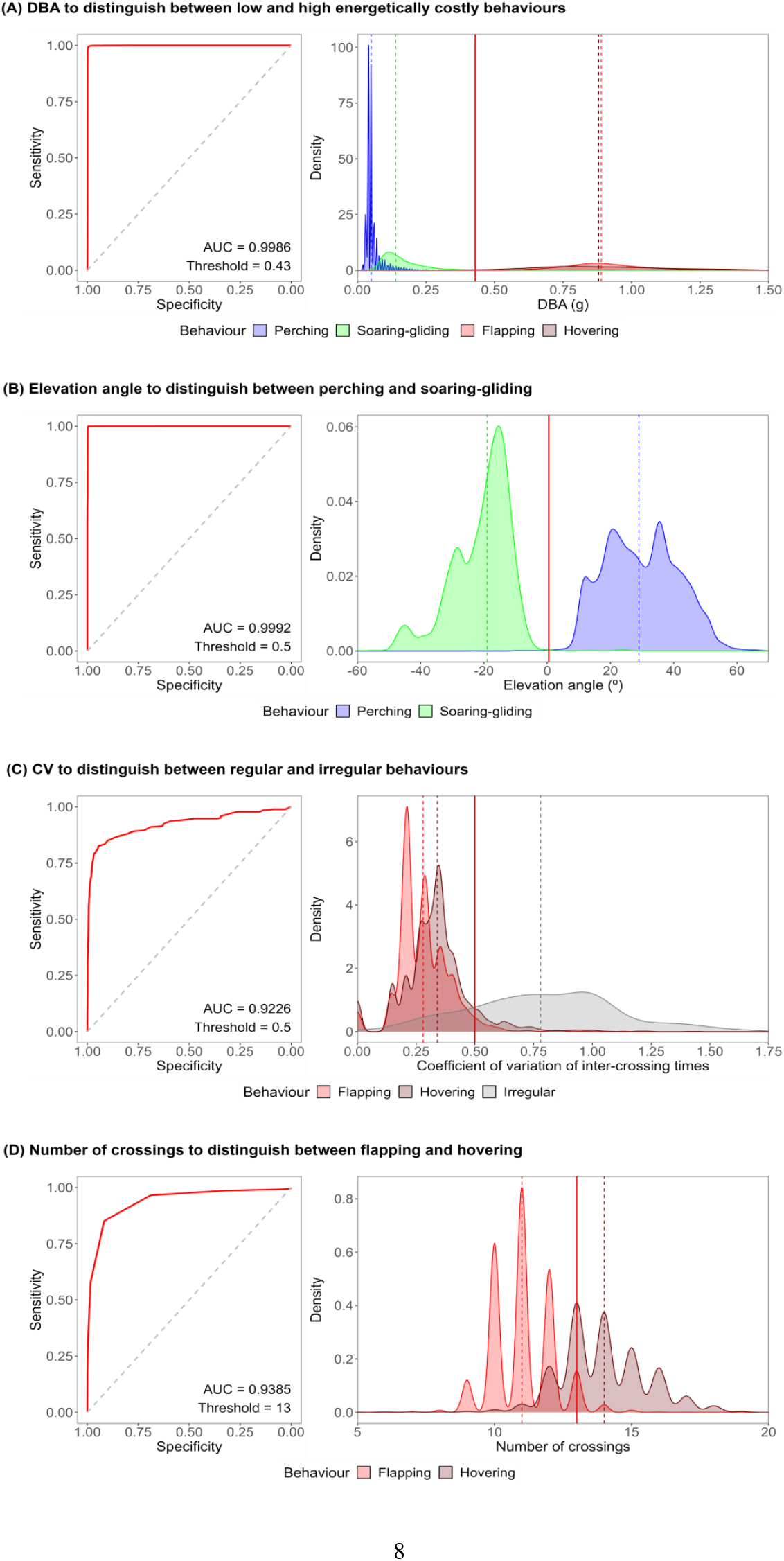
Selection of the threshold values based on receiver operating characteristic (ROC) analysis. Each panel shows the ROC curve and density plots for the parameter considered. The ROC curve was constructed by plotting the sensitivity (i.e. true positive rate) against the specificity (i.e. true negative rate). The area under the curve (AUC) measures the classifier quality of the parameter considered. The best possible threshold value was selected as the value providing the best trade-off between sensitivity and specificity (i.e. the closest ROC curve point to the upper left corner). In density plots, solid vertical red lines represent the selected threshold and dashed lines represent the median values of the behaviours involved.

As perching, soaring-gliding, flapping and hovering usually lasted a few seconds, we tested if the initial classification obtained by applying the decision tree to independent 1-s intervals could be improved by considering sequences of three or five consistent 1-s intervals. For this purpose, we tested 3 or 5-s width sliding windows and followed a ‘majority rule’: if the behaviour initially predicted for the 1-s interval at the centre of the window was preceded and followed by another behaviour, and both were the same, the central behaviour was re-classified to match the other two. For example, the sequence ‘Hovering-Flapping-Hovering’ was changed to ‘Hovering-Hovering-Hovering’ with the sliding window of 3-s (Table S1). Then, we assessed the accuracy of each of these three classifications (initial and done after the 3 or 5-s width sliding windows) by building a confusion matrix for each classification as compared to the ‘ground-truth’ classification obtained by manually labelling the various behaviours (Table 1).

### Identification of prey capture attempts

Sometimes, we observed the occurrence of distinctive high values of *a*_*z*_, typically represented by a single acceleration measurement over 25 in a 1-s interval (i.e. 1/25 s) (hereafter ‘spikes’, Fig. 5). In some cases, spikes were followed by straight flights to the nest box, suggesting that kestrels returned to feed the nestlings (Fig. 5A). Spikes were also preceded by sudden decreases in altitude and associated with upward and downward changes in body posture, as well as 1-s GPS sequences with zero instantaneous speed, suggesting that kestrel were handling prey (Fig. 5B). We therefore inferred that spikes were due to violent landings/strikes towards prey and considered them as signals of prey capture attempts, whether successful or not.

**Figure 5.**
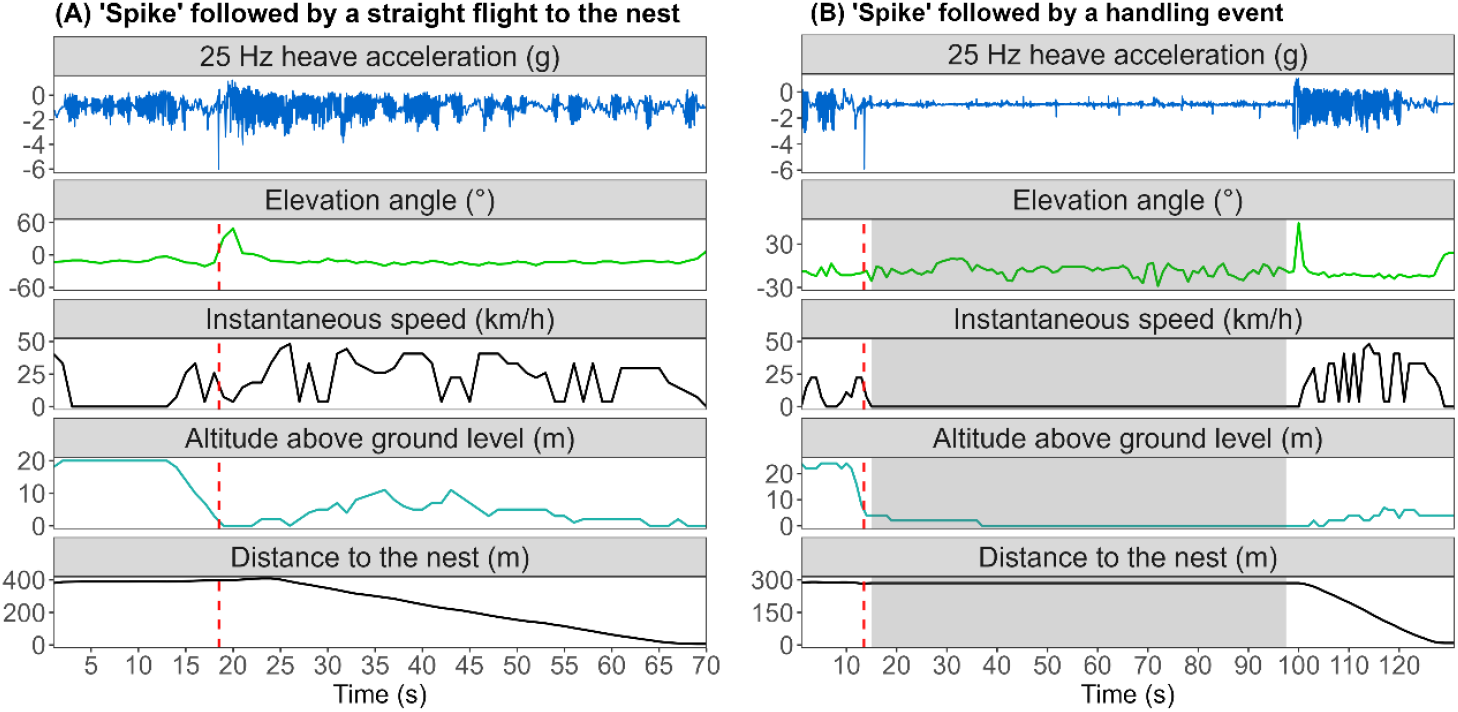
Successful prey capture attempt followed by a return to the nest (A) or by handling the prey item on the spot (B). Two examples that showed the changes in elevation angle, altitude above ground level and distance to the nest. Red dashed lines indicated the spikes and grey area in panel B the handling event. The altitude above ground level was estimated by subtracting the ground elevation in a 10-m resolution digital model elevation from the GPS-derived altitude above sea level.

To validate these inferences, we used the cameras monitoring the nest boxes to identify 70 prey delivery events and the associated prey capture attempts (62 events from 11 lesser kestrels and 8 events from two common kestrels). Video recordings confirmed that changes in elevation angles reflected prey handling because they also occurred when kestrels were dismembering prey to feed the nestlings inside the nest box. Using the the *a*_*z*_ values associated with manually labelled behaviours (Fig. S2A) and the spikes from validated prey capture attempts (Fig. S2B), we defined a spike as the occurrence of an absolute value of *a*_*z*_ higher than 3*g*. Additionally, we assumed that a spike preceded or followed by another spike, or spikes, within 30 seconds accounted for the same prey capture attempt. In these cases, the spike with the highest value of *a*_*z*_ was considered as the 1-s interval reference for the event. To assess the hunting method used by the kestrel, we estimated the time percentage of perching behaviour within the 30-s interval that preceded the spike. It was considered perch-hunting or flight-hunting depending on whether the percentage was higher or lower than 50% (i.e. 15 s).

## Results

The three automatic classifications reached a very high accuracy, larger than 97%. The best accuracy was obtained with the 3-s width sliding window, but the gain in accuracy obtained in this way was marginal (Table 1).

We successfully identified the 1-s interval of reference for 97% of the validated prey capture attempts (68 events). The two remaining events could not be identified as their spikes were equal to 3*g* (see Fig. S2B). Regarding the full dataset, we detected 759 prey capture attempts: 709 from lesser kestrels (5.7 attempts per tracking hour) and 50 from common kestrels (1.5 attempts per tracking hour). Among lesser kestrels, 63 attempts (9%) were classified as perch-hunting and 646 (91%) as flight-hunting, while for common kestrels, 12 attempts (24%) were classified as perch-hunting and 38 (76%) as flight-hunting.

## Discussion

Since the first studies on animal behaviour using tri-axial high-frequency accelerometers were published 25 years ago (reviewed in Brown et al. 2013, Wilson et al. 2020), a variety of methods have been developed to identify/classify behaviours, including histogram analysis (Collins et al. 2015), classification and regression trees (Shamoun-Baranes et al. 2012), *k*-means cluster analysis (Bidder et al. 2014), hidden Markov models (Leos-Barajas et al. 2017), artificial neural networks (Nathan et al. 2012, Jeantet et al. 2021) and random forests (Pagano et al. 2017, Jeantet et al. 2020). On the other hand, simpler methods often use threshold values of parameters derived from raw acceleration data that are typically estimated either by subjective inspection (Yoda et al. 2004, Holland et al. 2009) or by comparison with ground-truth data (e.g. video recordings: Halsey et al. 2009, field observations: Shamoun-Baranes et al. 2012, or both: Moreau et al. 2009, Fluhr et al. 2021).

Our study, based on a simple threshold-based approach, demonstrated that kestrels’ behaviours can be automatically predicted with a very high accuracy (>97%) by using a simple decision tree based on four parameters derived from the data acquired by a tri-axial high-frequency accelerometer. Although we did not directly compare our method with others, the accuracy reached by our approach is relatively higher than most other studies relying on more sophisticated classifications(e.g. 91% for seven behaviours of griffon vultures *Gyps fulvus*, Nathan et al. 2012; 77% for five behaviours of imperial cormorants *Leucocarbo atriceps*, Bidder et al. 2014; 92% for five behaviours of white stork *Ciconia ciconia*, Yu et al. 2021; 91-96% for five behaviours of green turtle *Chelonia mydas*, Jeantet et al. 2020, 2021). Our study also showed that kestrels’ prey capture attempts can be objectively identified from diagnostic acceleration signatures (the so-called ‘spikes’). This provides an exciting opportunity to investigate the hunting behaviour of aerial predators with no need of direct observation. Most classification models based on machine learning tend to be fed with a large number of parameters that are not necessarily biologically relevant such as raw acceleration values. For instance, in a sample of 15 studies, Patterson et al. (2019) showed that 12 parameters were used on average to classify behaviours. In contrast, our aim was to develop a simple method involving an explicit decision tree based on a few biologically relevant parameters that are easy to interpret. In particular, the activity level (DBA) and the elevation angle made it possible, in our study species, to distinguish between perching and soaring-gliding flight with a very high precision (99%, Table 1). In line with other studies, these two parameters were useful to identify grazing in cows *Bos taurus* (Rayas-Amor et al. 2017), running in squirrels *Tamiasciurus hudsonicus* (Studd et al. 2019) or swimming in seabirds (Patterson et al. 2019).

Using a simple decision tree should also make our method much more easily applied to a large number of aerial species, as this would only require to change the threshold values in the decision tree. The accuracy may be lower in comparison with machine learning methods (Moreau et al. 2009, Shamoun-Baranes et al. 2012), but this is not necessarily the case (Studd et al. 2019; Fluhr et al. 2021). A potential limitation in a number of studies involving an explicit decision tree is the way the various thresholds were determined. In this paper, we showed that this can be done objectively based on a ROC analysis.

The approach of building an explicit decision tree is nevertheless subject to some drawbacks when different behaviours generate similar acceleration signatures and can only be distinguished based on a large number of parameters (Moreau et al. 2009, Shamoun-Baranes et al. 2012). For these cases, learning machine approaches are certainly the best choice as they are able to manage simultaneously large amounts of data of different natures (Jeantet et al. 2020, 2021). Thus, in our study, dealing with the number of times that heave acceleration crosses –1*g* to distinguish between flapping and hovering flights was tricky, with the latter behaviour being the least precisely identified (ca. 85%, Table 1). Indeed, under suitable wind conditions (ca. 8 m/s, Videler and Groenewold 1991), kestrels can hover with reduced wingbeat frequency and thus an overlap between these two flight modes may occur (Fig. 4D). Crucially, knowledge of the target species and the target behaviours may allow for the inclusion of a biological context to further enhance the classification. Considering that bouts of behaviours last for a few seconds (Village 1990, Tella et al. 1998, Vlachos et al. 2003, Hernández-Pliego et al. 2017b), we improved the original precision of flapping and hovering by approximately 2% with the application of a ‘majority rule’ within sliding windows of varying time lengths (Table 1). Although the use of GPS instantaneous speed would have easily solved this issue as hovering is characterised by zero speed, we aimed at providing an approach based exclusively on acceleration data that could be readily applied to larger datasets without paired high-frequency GPS data, which ultimately leads to reduced sampling durations due to its high energy consumption (Bennison et al. 2018). Furthermore, kestrels are known to show short irregular movements in between bouts of flapping, hovering and/or soaring-gliding flights (Rijnsdorp et al. 1981, King and Cowlishaw 2009), and only a few methods such as Hidden Markov models can directly model these expected transitions (Leos-Barajas et al. 2017). However, as mentioned earlier, there are often simpler alternatives. By including the CV of the delay between successive wingbeats (using the acceleration along the heave axis as a proxy) in the classification model, we ensured an accurate identification of irregular moves and improved the overall accuracy. Finally, it is worth noting that using high-frequency GPS data as ground-truth constitutes a valuable solution when the most common sources of validation are unobtainable or unlikely to represent the full range of predefined behaviours, as previously noted by Pagano et al. (2017) and Patterson et al. (2019). However, the manual task of labelling behaviours is labour intensive and potentially discouraging. The development of an interactive web app makes the process easier and faster (Fig. S1).

The remote detection of very quick events such as prey capture attempts remains a challenge (Clermont et al. 2021). This is the case for studies on aerial predators like kestrels (Rijnsdorp et al. 1981, Masman et al. 1988, Village 1990, Zank and Kemp 1996, Tella et al. 1998, Vlachos et al. 2003). Hernández-Pliego et al. (2017b), who studied the foraging behaviour of lesser kestrels with paired GPS and acceleration data, were not able to identify prey capture attempts. Our success here resulted from considering brief and strong changes in acceleration data that corresponded to dives or strikes towards prey on the ground. As in previous studies (Masman 1988, Village 1990, Vlachos et al. 2003), we showed that lesser and common kestrels use more frequently the flight-hunting strategy (i.e. hovering) when provisioning the nestlings because it is more time-efficient for finding prey than perch-hunting, although more energy-consuming (Masman 1988, Hernández-Pliego et al. 2017b). Taking into account that our spike-based method should only be considered as a first step in the use of acceleration data to objectively study the hunting behaviour of aerial predators, we would like to mention a couple of limitations. Without additional information, the hunting success can only be poorly assessed based on the possible detection of a handling event following a successful attempt. Based on GPS data and video recordings at the nest, we observed that prey can be captured without generating a clear handling signature. Thus, dives that do not generate spikes (e.g. two out of 70 attempts in our sample obtained with video recording, Fig. S2B) and successful self-feeding events that require extremely short handling times or none at all (e.g. small insects, Sonerud et al. 2014), cannot be identified. Further research should seek to refine our spike-based method in order to be able to detect successful hunting events with devices requiring less power than GPS loggers such as accelerometers and gyrometers.

## Acknowledgments

We thank M. Baena, C. Marfil and P. Aycart for their help during the fieldwork. We especially thank M. J. Smith for his help with cameras and M. Vázquez for his help in the capture, ringing and tagging. This work was supported by “KESTRELS-MOVE” (CGL2016-79249-P) (AEI/FEDER, UE), “MERCURIO” (PID2020-115793GB) (AEI/FEDER, EU), “SUMHAL” (LIFEWATCH-2019-09-CSIC-13, POPE 2014-2020) (MICINN, EU) and “SIMMRE” project (BIOD22_00033_3_PPCB) (PCBIO Junta de Andalucía, PRTR/NextGeneration, EU). DGS was financially supported by the Spanish Ministry of Education, Culture and Sports for university teacher training (FPU) (FPU17/04342). We thank ICTS-RBD for giving logistic and technical support in Doñana.

## Appendix

**Table S1.**
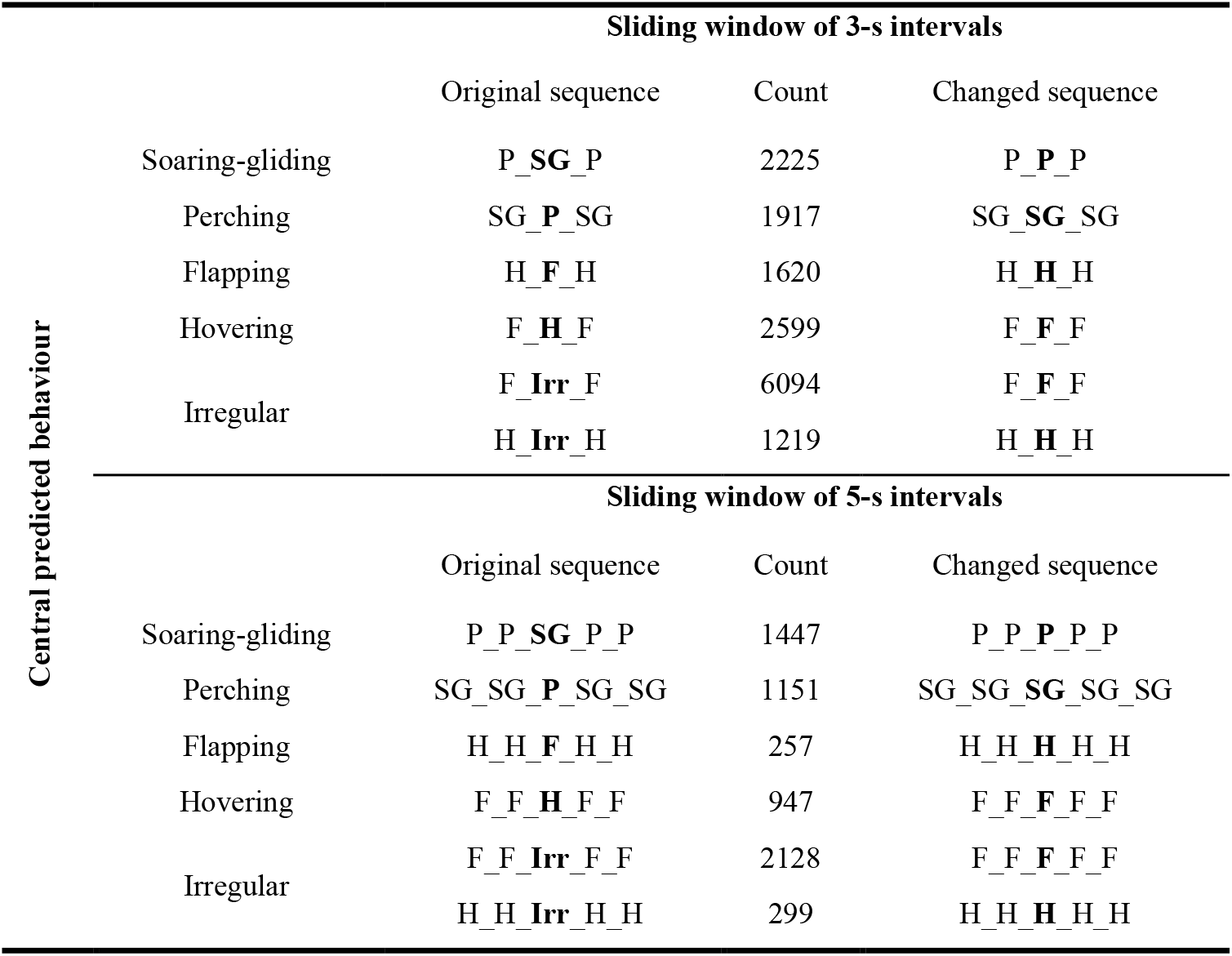
Changes made according to the majority rule (3-s and 5-s intervals). For each sliding window, the original sequence, the number of cases and the resulting sequence after being changed are indicated. P represented perching, SG soaring-gliding, F flapping, H hovering and Irr irregular

**Figure S1.**
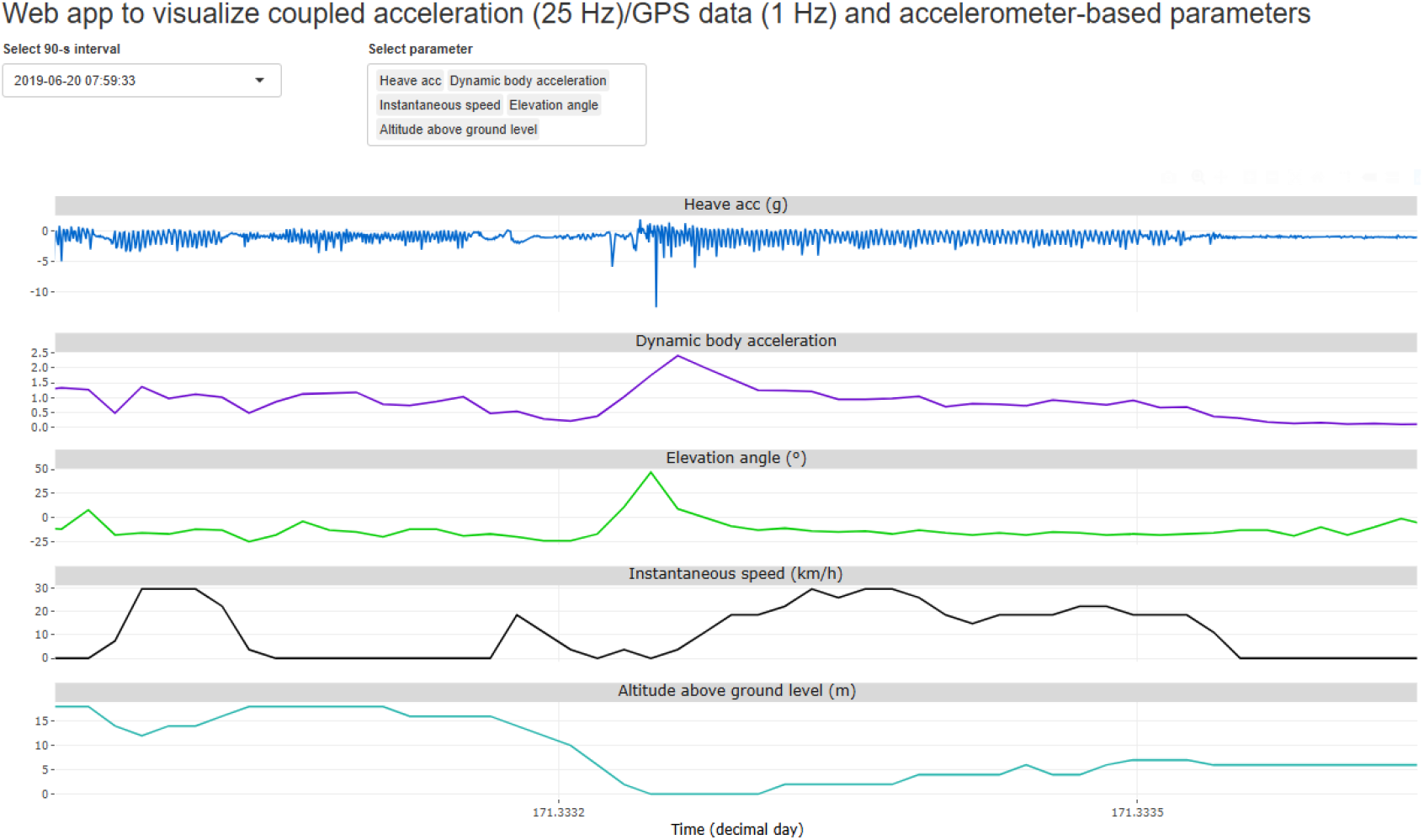
Interactive web app created with the R package ‘Shiny’ (Chang et al. 2022). It combined raw acceleration data at 25 Hz (surge, sway and heave axes), parameters at 1-second interval derived from acceleration data (dynamic body acceleration, elevation angle, the number of crossings and the coefficient of variation of inter-crossing times) and GPS information (altitude above ground level and instantaneous speed). The altitude above ground level was estimated by subtracting the ground elevation in a 10-m resolution digital model elevation from the GPS measured altitude above sea level.

https://danielgarciasilveira.shinyapps.io/ShinyApp/

**Figure S2.**
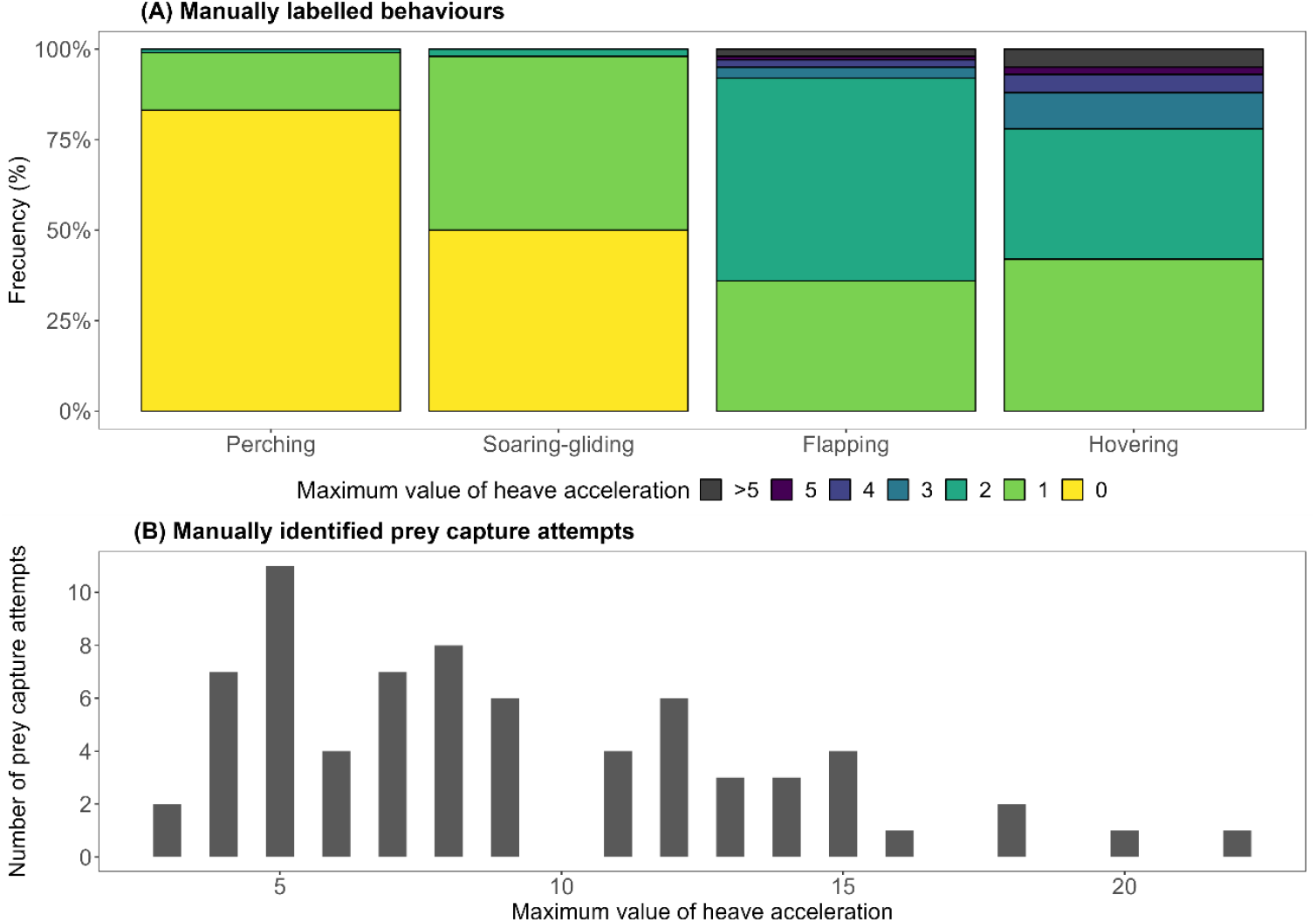
Maximum values of heave acceleration. Panel A indicated the values associated with each manually labelled behaviour, for example soaring-gliding flight was mainly characterised by values of 0 and 1*g*, and flapping flight by 1*g* and 2*g*. Panel B indicated the number of prey capture attempts per value of heave acceleration, for example seven attempts had spikes of 4*g* and 11 events showed spikes of 5*g*.

## Notes

### Competing Interest Statement

The authors have declared no competing interest.

